# Distinct neuromodulatory effects of endogenous orexin and dynorphin corelease on projection-defined ventral tegmental dopamine neurons

**DOI:** 10.1101/2024.08.01.606179

**Authors:** Aida Mohammadkhani, Min Qiao, Stephanie L. Borgland

## Abstract

Dopamine (DA) neurons in the ventral tegmental area (VTA) respond to motivationally relevant cues and circuit-specific signaling drives different aspects of motivated behavior. Orexins (ox; also known as hypocretin) and dynorphin (dyn) are co-expressed lateral hypothalamic (LH) neuropeptides that project to the VTA. These peptides have opposing effects on the firing activity of VTA^DA^ neurons via orexin 1 (Ox1R) or kappa opioid (KOR) receptors, respectively. Given that Ox1R activation increases VTA^DA^ firing, and KOR decreases firing, it is unclear how the co-released peptides contribute to the net activity of DA neurons. We tested if optical stimulation of LH neuromodulates VTA^DA^ neuronal activity via peptide release and if the effects of optically driven LH_ox/dyn_ release segregates based on VTA^DA^ projection targets including the basolateral amygdala (BLA) or the lateral or medial shell of the nucleus accumbens (lAcbSh, mAchSh). Using a combination of circuit tracing, optogenetics, and patch clamp electrophysiology in male and female orexin^cre^ mice we showed a diverse response of LH optical stimulation on VTA^DA^ neuronal firing, that are not mediated by fast transmitter release and are blocked by antagonists to KOR and Ox1R signaling. Additionally, where optical stimulation of LH_ox/dyn_ inputs in the VTA inhibited firing of the majority of BLA projecting VTA^DA^ neurons, optical stimulation of LH inputs in the VTA bidirectionally affects firing of either lAcbSh or mAchSh projecting VTA^DA^ neurons. These findings indicate that LH_ox/dyn_ corelease may influence the output of the VTA by balancing ensembles of neurons within each population which contribute to different aspects of reward seeking.

**Significance Statement:** The mesolimbic dopamine (DA) system is known to play a crucial role in motivation and reward-learning and receives neuromodulatory input from the lateral hypothalamus (LH). We show that optical stimulation of the orexin-containing LH input in the VTA releases both orexin and dynorphin to bidirectionally alter VTA^DA^ firing. Furthermore, orexin and dynorphin differentially modulate firing of DA inputs to the basolateral amygdala, whereby dynorphin predominates, or to the nucleus accumbens which is sensitive to both neuromodulators. Our findings contribute to a more comprehensive understanding of the neuromodulatory effects of coreleased LH orexin and dynorphin on the VTA^DA^ system.

## Introduction

Reward and reinforcement processes drive motivated behaviors and are essential for survival and are guided by the ventral tegmental area (VTA) (Berridge and Robinson, 1998; Bass et al., 2013; Wang et al., 2015; Morales and Margolis, 2017). Dopamine (DA) neurons in the VTA integrate multiple inputs to encode signals influencing motivated behaviors through diverse projections, including projections to nucleus accumbens (NAc) (Beier et al., 2019) and amygdala subnuclei, such as the basolateral amygdala (BLA) (Lammel et al., 2008; Morel et al., 2022). VTA^DA^ projections to the NAc and BLA are anatomically segregated, such that DA neurons do not send collaterals to both regions (Beier et al. 2015; Baimel et al. 2017).

The VTA also receives input from orexin (ox, also known as hypocretin) neurons of the lateral hypothalamus (LH). LH_ox_-containing fibers project to the VTA and make close appositions to VTA^DA^ neurons (Peyron et al., 1998; Fadel and Deutch, 2002; Baldo et al., 2003). While retrograde labeling from the VTA reveals a significant LH_ox_ input to the VTA (González et al., 2012), other studies have indicated that LH_ox_ neurons synapse onto about 5% of VTA^DA^ and GABA neurons, even though there are many orexin-containing dense core vesicles within the VTA (Balcita-Pedicino and Sesack, 2007). Orexin interacts with orexin 1 (Ox1R) and 2 (Ox2R) receptors expressed in the VTA (Marcus et al., 2001; Narita et al., 2006), which are thought to primarily couple to Gq proteins as activation of Ox1R increases intracellular calcium (Uramura et al., 2001), endocannabinoids (Tung et al., 2016), and increases firing of VTA^DA^ neurons (Korotkova et al., 2003; Baimel et al., 2017). Orexin microinjected in the VTA can increase DA release in the NAc (Vittoz and Berridge, 2006; España et al., 2011). This is consistent with the findings that optogenetic activation of the LH orexin input to the VTA promotes DA release in the NAc. This occurs under phasic release conditions, likely resulting from the stimulation of glutamatergic afferents to the VTA (Thomas et al., 2021). Thus, orexin strengthens the activity and output of dopaminergic neurons that project to the NAc. As such, orexin action in the VTA is linked with motivational processes (Tyree and de Lecea, 2017). In particular, orexin signaling in the VTA increases motivation for highly salient rewards such as addictive drugs or energy dense foods (Borgland et al., 2009; Thompson and Borgland, 2011).

Orexins are colocalized with dynorphin in approximately 95% of neurons (Chou et al., 2001; Li and van den Pol, 2006). These peptides are also co-expressed within dense core vesicles (Muschamp et al., 2014), suggesting that they are co-released. Dynorphin is the endogenous ligand of Gi/o coupled kappa opioid receptors (KORs), which are expressed on somatodendrites of VTA^DA^ neurons (Abraham et al., 2022). Activation of KORs in the VTA inhibit firing (Margolis et al., 2003, 2006; Ford et al., 2006), suppress excitatory input to VTA^DA^ neurons (Margolis et al., 2005), and reduce DA release in the NAc (Robble et al., 2020). Thus, given the opposing action of orexin and dynorphin, how does co-release of orexin and dynorphin alter the activity of VTA^DA^ neurons? When both orexin and dynorphin are applied to VTA^DA^ neurons that are responsive to either agonist individually, there is no net effect on firing rate, suggesting that the opposing effects of each peptide effectively cancel one another out (Muschamp et al., 2014). However, few dopaminergic neurons are simultaneously responsive to both KOR and Ox1R activation (Muschamp et al., 2014). Electrical stimulation of miniature brain slices containing LH orexin neurons released dynorphin, measured with an enzyme linked immunosorbent assay (ELISA) using a dynorphin antibody (Li and van den Pol, 2006). Exogenous application of orexin or dynorphin to the LH (Li and van den Pol, 2006) or the VTA (Baimel et al., 2017) produces opposing effects on firing. However, it is unknown how subpopulations of VTA^DA^ neurons that participate in distinct circuits differentially respond to the neuromodulatory influence of LH_ox/dyn_ neurons. We combined circuit tracing, optogenetics and whole cell patch clamp recordings to investigate 1) if the LH_ox/dyn_ input to the VTA can co-release orexin and dynorphin to modulate VTA^DA^ neuronal firing, and 2) if VTA^DA^ neuronal projections to the lateral shell of the NAc (lAcbSh), the medial shell of the NAc (mAcbSh), or the BLA are differentially modulated by LH_ox/dyn_ input. We hypothesized that photoactivating the LH_ox/dyn_ input to the VTA induces activation or inhibition of distinct projection defined subpopulations of VTA^DA^ neurons via orexin or dynorphin respectively.

## Materials and Methods

### Subjects

Adult male and female orexin^cre^ mice (post-natal day 60-90) were originally obtained from the Yamanaka lab at the University of Tokyo (Inutsuka et al., 2014) and bred locally at the University of Calgary Clara Christie Center for Mouse Genomics. Mice were group-housed (3-5 same sex per cage) with *ad libitum* access to water and food. Mice were housed in ventilated cages in a temperature (21 ± 2°C) and humidity-controlled (30-40%) room on a 12h reverse light/dark cycle (lights on at 08.00 AM). Experiments were performed during the animal’s light cycle. All experimental procedures adhered to ethical guidelines established by the Canadian Council for Animal Care and animal use protocols approved by the University of Calgary Animal Care and Use Committee (AC21-0034).

### Surgical procedures

All orexin^cre^ mice (mice expressing the cre recombinase in cells expressing the pre-pro-orexin peptide (Inutsuka et al., 2014) received bilateral infusions of either channelrhodopsin (‘ChR2’) (AAV2/8-EF1a-DIO-hChR2(H134R)-mCherry; Neurophotonics, Centre de Recherche CERVO, Quebec City, QC, Canada) or control (‘mCherry’) (AAV2/8-hSyn-DIO-mCherry; Neurophotonics) virus. Mice were anaesthetised with isoflurane gas (5% for induction; 1-2% for maintenance) and secured in a stereotaxic frame (David Kopf Instruments, Tujunga, CA, USA). All measurements were made relative to bregma for viral infusions. Viral injections were performed using a microinjector (Nano-inject II; Drummond Scientific Company, Broomall, PA, USA). Each mouse received 6 infusions into the LH (100 nl per infusion; 23.1 nl/s), 3 in each hemisphere (anterior-posterior (AP) -1.35; mediolateral (ML) ± 0.9; dorsoventral (DV) -5.2, -.5,1, -5.0) for a total of 300 nl per hemisphere. After each infusion, the microinjector was left in place for 3 min to allow diffusion of virus away from the needle tip. After all infusions were complete in one hemisphere, the microinjector was left in place for an addition 5 min, 500 μm dorsal of the final injection location to allow diffusion of the virus through the brain tissue. Red Retrobeads (max excitation at 530 nm/ max emission at 590 nm) (200 nL; Lumafluor Inc., Naples, FL) were infused bilaterally into the lAcbSh (from bregma: anteroposterior +1.425 mm; mediolateral, ± 1.75 mm; dorsoventral, −4.25 mm), the mAcbSh (from bregma: anteroposterior: +1.65 mm; mediolateral, ± 0.5 mm; dorsoventral, -4.6), or the BLA (from bregma: anteroposterior -1.0 mm; mediolateral, ± 3.1 mm; dorsoventral, -5.3 mm). Injection sites were confirmed in all animals by preparing coronal section of the lAcbSh or mAcbSh and in horizontal sections of the BLA. All mice received pre- and postoperative analgesic (Meloxicam 5 mg/kg, subcutaneous) and were returned to their home cages and allowed to recover for 6-8 weeks prior to further experimental procedures. The location of the virus expression was performed posthoc.

### Electrophysiology

All electrophysiological recordings were performed in slice preparations from adult male and female orexin^cre^ mice 6 weeks after receiving the cre-dependent viral vector containing ChR2. Briefly, mice were anaesthetized with isoflurane and transcardially perfused with an ice-cold N-methyl-d-glucamine (NMDG) solution containing (in mm): 93 NMDG, 2.5 KCl, 1.2 NaH2PO4.2H2O, 30 NaHCO3, 20 Hepes, 25 D-glucose, 5 sodium ascorbate, 3 sodium pyruvate, 2 thiourea, 10 MgSO4.7H2O and 0.5 CaCl2.2H2O and saturated with 95% O2–5% CO2. Mice were then decapitated, and brains were extracted. Sections containing the Lumifluor injection site were confirmed visually. Horizontal sections (250 μm) containing the VTA were cut in NMDG solution using a vibratome (VT1200, Leica Microsystems, Nussloch, Germany). Slices were then incubated in NMDG solution (32°C) saturated with 95% O2–5% CO2 for 10 min before being transferred to a holding chamber containing artificial cerebrospinal fluid (ACSF), (in mM): 126 NaCl, 1.6 KCl, 1.1 NaH2PO4, 1.4 MgCl2, 2.4 CaCl2, 26 NaHCO3 and 11 glucose (32–34°C) equilibrated with 95% O2–5% CO2 for at least 45 min before recording. Slices were transferred to a recording chamber on an upright microscope (Olympus BX51WI) and continuously superfused with ACSF (2 mL/min, 34 °C). Cells were visualized on an upright microscope using “Dodt-type” gradient contrast infrared optics and whole-cell recordings were made using a MultiClamp 700B amplifier (Axon Instruments, Molecular Devices). Recording electrodes (3-5 MΩ) for measuring firing rates were filled with potassium-D-gluconate internal solution (in mM): 130 potassium-D-gluconate, 4 MgCl2, 10 HEPES, 0.5 EGTA, 10 sodium creatine phosphate, 3.4 Mg-ATP and 0.3 Na2GTP and 0.2% biocytin.

After breaking into the cell, hyperpolarization-activated cation currents (Ih) were recorded in voltage-clamp mode using a voltage step to -130 mV to DA neurons voltage-clamped at -70 mV. Ih was determined as the change in current between ∼30 ms and 248 ms after the voltage step was applied. Because most DA neurons ceased firing within 5 min of recording, current-step induced firing was used for all experiments. For current-step experiments, the membrane potential for each neuron was set to - 60 mV by DC injection via the patch amplifier and a series of 5 current pulses (250 ms in duration, 5-25 pA apart, adjusted for each cell) were applied every 45 s, where the minimum current amplitude was set for each cell so that the first pulse was subthreshold and did not yield firing. From this series of current steps, we then selected a current step that yielded 3-5 action potentials during the baseline period and used that step for the analysis of peptide effects, as described previously (Baimel et al., 2017).

We optically stimulated LH_ox/dyn_ inputs at 30 Hz over a range of durations (10, 20 or 30 s) from a light-emitting diode (LED) blue light source (470nm) directly delivered the light path through the Olympus 40X water immersion lens. Once a maximal stimulation was determined, we continued with 30 Hz, 30s stimulation for further experiments. SB334867 and NorBNI (Ox1R antagonist, KOR antagonist; Tocris; 1 µM) were both dissolved in 100% DMSO stock solutions and then diluted to their final concentration containing 0.001% DMSO in ACSF and bath applied to slices.

### Analysis of action potential firing

Firing data for all neurons was analyzed with the MiniAnalysis program (Synaptosoft). Optical stimulation-induced changes in firing are expressed as a percentage of baseline. For analysis of the time courses, the firing rate pre- and post-optical stimulation was normalized to the average of the 10 min baseline firing rate. For analyses of effect sizes, the last 2 min prior to the optical stimulations or at the end of the recordings were averaged. To distinguish responders showing a decrease or increase from non-responders, we used a criterion of a 20% change in firing rate from the baseline. Responses from neurons of male and female mice were analyzed together. Our pilot data indicated that there were not sufficiently large sex differences in electrophysiological responses and therefore mice were grouped together as these studies were not sufficiently powered to test for sex differences.

All statistical analyses were performed in GraphPad Prism 9.4.1 (GraphPad, US). Paired t-test were used to compare before and after drug applications. In the case where 3 or more timepoints were compared, a repeated measures ANOVA was used. In all electrophysiology experiments, sample size is expressed as N/n where “N” refers to the number of cells recorded from “n” animals. Recordings of male and female mice were grouped together due to limited availability of the mice. All cells were then averaged together and presented as mean ± SEM with individual values overlayed. All significance was set at P<0.05. The levels of significance are indicated as follows: ****p<0.0001, ***p< 0.001, **p < 0.01, *p < 0.05.

### Immunohistochemistry and confocal microscopy

The VTA is composed of a heterogeneous collection of cell types, distinguished in part by neurotransmitter content. To determine whether recorded neurons are indeed DAergic, we filled neurons with biocytin while recording and subsequently processed slices for tyrosine hydroxylase (TH). Brain slices from patch-clamp recordings in orexin^cre^ mice were fixed overnight in cold 4% paraformaldehyde and then stored in phosphate-buffered saline until processing. Sections were then blocked in 10% normal goat serum and incubated with mouse anti-TH (1:1000) for 24 hours at room temperature. Alexa Fluor 488 goat anti-mouse (1:400) and Dylight 594 conjugated streptavidin were applied to identify DA neurons tagged with biocytin. Slices were mounted with Fluoromount.

To check for co-localization of ChR2 expression and orexin in LH, mice were deeply anesthetized with isoflurane and transcardially perfused with phosphate buffered saline (PBS) and then with 4% paraformaldehyde (PFA). Brains were dissected and post-fixed in 4% PFA at 4°C overnight, then switched to 30% sucrose. Coronal frozen sections were cut at 30 µm using a cryostat. 10% goat serum was applied to block non-specific binding for 1 h. Sections were then incubated with primary antibody rabbit anti-orexin 1:500 (Phoenix pharmaceuticals, H-003-30) and chicken red fluorescent protein (RFP, to enhance mCherry reporter expression) 1:2000 (Rockland, 600-901-379) in 1% BSA for 1 h at room temperature followed by incubation with secondary antibody Alexa Fluor 488 goat anti-rabbit and Alexa Fluor 594 goat anti chicken 1:400 for 1 h.

To check for the colocalization of retrobeads, biocytin and TH in VTA patched slices, 10% goat serum was applied to block non-specific binding for 1 h. Sections were then incubated with primary antibody mouse anti-TH (1:1000) in 1% BSA for 24 h at room temperature followed by incubation with secondary antibody Alexa Fluor 647 goat anti mouse (1:400) and Alexa fluor 488 streptavidin (1:200) for 2 hours.

All images were obtained at 10x on an Olympus scanner microscope (Olympus Canada, Ontario, Canada) and at 20x on a Nikon Eclipse C1si confocal microscope (Nikon Canada Inc. Ontario, Canada). Cell count in LH was quantified with ImageJ at 20x.

## Results

### Optical stimulation of LH_ox/dyn_ inputs to VTA^DA^ neurons bidirectionally modulates firing

To study the effect of LH photoactivation on VTA^DA^ neuronal activity we injected AAV2/8-EF1a-DIO-hChR2(H134R)-mCherry or control (‘mCherry’) (AAV2/8-hSyn-DIO-mCherry) virus into the LH of orexin^cre^ mice (Fig. 1 A, B, C). 81 ± 12% of LH orexin neurons expressed the reporter for ChR2 and 91 ± 3% of ChR2-expressing neurons were orexin+. Because dynorphin is expressed in orexin-containing neurons, and no other neurons in the LH, this manipulation is specific to the LH_ox/dyn_ population. We recorded the response of VTA^DA^ neurons, identified by electrical characteristics and posthoc TH staining, after optical stimulation of LH_ox/dyn_ inputs in the VTA, using whole cell patch clamp electrophysiological recordings (Fig 1D-F). Following the baseline recording, we applied optical stimulation at a frequency of 30 Hz with increasing duration, which has been shown to effectively modulate the firing of VTA^DA^ neurons previously (Thomas et al., 2022). If an increase in firing frequency occurred in response to optical stimulation, we applied the Ox1R antagonist, SB334867 (1 μM, 15 min), as orexin is known to increase firing of VTA^DA^ neurons (Korotkova et al., 2003; Baimel et al., 2017; Thomas et al., 2022). Conversely, if a decrease in firing occurred, we administered the KOR antagonist, NorBNI (1 μM, 15 min), as dynorphin is known to decrease firing of VTA^DA^ neurons (Margolis et al., 2003; Baimel et al., 2017). We did not apply an antagonist if no change in firing occurred. Optical stimulation of LH inputs to VTA^DA^ neurons did not alter firing rate in mCherry mice (baseline, 100 ± 0.03%; 30Hz,10s: 106 ± 4%; 30Hz,20s: 114 ± 4%, 30Hz,30s: 112 ± 5%; n/N=17/13; Fig. 1G), suggesting that the increasing duration of the 473 nM LED does not alter firing on its own. In contrast, optical stimulation of LH inputs to VTA^DA^ neurons of ChR2 expressing mice produced diverse responses. In 37% of VTA^DA^ neurons, there was a duration dependent increase in firing (baseline: 104% ± 4%; 30 Hz, 10 s: 121 ± 10%; 30 Hz, 20 s: 143 ± 10%; 30 Hz,30 s: 151 ± 10%; SB34867: 131 ± 5%; n/N=7/5; Fig. 1H, I-K). Increased firing was significantly different from baseline at 30 Hz, 20s (p = 0.035) and 30 Hz, 30s (p = 0.016), but not after application of SB334867 (p = 0.57), (RM one way ANOVA: F(2.23,13.4)=6.52, p=0.0091; Fig. 1J). In 47% of VTA^DA^ neurons, there was a duration dependent decrease in firing (baseline: 99 ± 0.5%; 30 Hz,10 s: 73 ± 6%; 30 Hz, 20 s: 59 ± 7%; 30 Hz,30 s: 56 ± 4%; NorBNI: 92 ± 16%; n/N = 9/5; Fig. 1H). Decreased firing was significantly different from baseline at 30 Hz, 10 s (p < 0.0001), 30 Hz, 20 s (p < 0.0001), and 30 Hz, 30 s (p < 0.0001), but not after application of NorBNI (p = 0.57), (RM one way ANOVA: F(4,32)=19.26, p< 0.0001; Fig. 1L, M). A subset of VTA^DA^ neurons (16%) showed no difference in firing (Fig. 1H,I).

**Figure 1.**
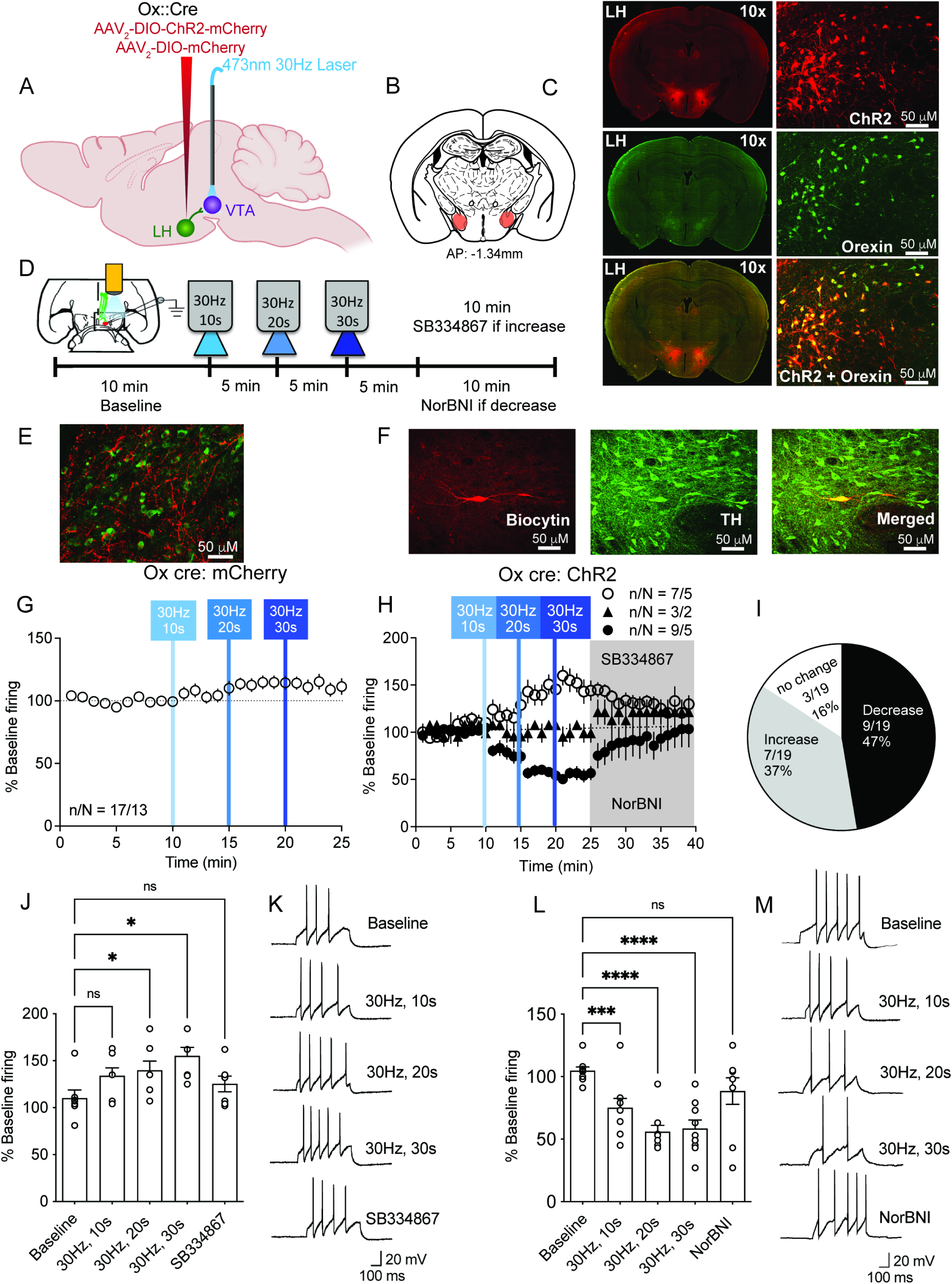
Optical stimulation of LH _ox/dyn_ inputs in the VTA produce a diverse response on firing rate of VTA^DA^ neurons. **A)** Schematic of viral strategy. Orexin^cre^ mice were infused with either channelrhodopsin (AAV2/8-EF1a-DIO-hChR2(H134R)-mCherry; ‘ChR2’) or control virus (AAV2/8-hSyn-DIO-mCherry; ‘mCherry’). (LH: lateral hypothalamus; VTA: ventral tegmental area. **B)** Coronal brain sections mapping viral infection areas within the LH (-1.34mm in anterior-posterior plane relative to Bregma). **C)** Immunohistochemical demonstration of cre-dependent expression of ChR2-mCherry (red, upper), orexin (green, middle), and a merged image of orexin and mCherry (ChR2 reporter) within the LH (lower). Scale bar, 50 µm. **D)** Experimental timeline of electrophysiology experiments with repeated optical stimulation. **E)** Example image of LH fibers expressing ChR2 within the VTA of orexin^cre^ mice (merged image of VTA^DA^ neuron labeled with tyrosine hydroxylase (TH, green) and LH fibers expressing ChR2 (mCherry reporter, red) **F)** VTA^DA^ neuronal identification by co-labeling of biocytin (red, left) and TH (green, middle) or the merged image of VTA DA neuron co-labeled with TH and biocytin (right). Scale bar, 25 µm. **G)** Time course of evoked firing before and after 30 Hz photoactivation of LH_ox/dyn_ inputs in VTA DA neurons (10s, 20s, 30s) of mCherry expressing control mice. **H)** Time course of evoked firing before and after 30 Hz photoactivation of LH_ox/dyn_ inputs on VTA^DA^ neurons (10s, 20s, 30s) of ChR2-expressing mice. **I)** Distribution of responses of VTA^DA^ neurons to LH photostimulation. **J)** Bar graph of averaged firing responses (% baseline) of the 2 min prior to the next optical stimulation for VTA^DA^ neurons that increased their firing to optical stimulation of LH_ox/dyn_ inputs. Individual responses are overlayed. **K)** Sample traces of evoked action potentials that increased frequency before and after optical stimulation of LH_ox/dyn_ inputs. **L)** Bar graph of averaged firing responses (% baseline) from 2 min prior to the next optical stimulation for VTA^DA^ neurons that decreased their firing to optical stimulation of LH_ox/dyn_ inputs. **M)** Sample traces of evoked action potentials that decreased frequency before and after optical stimulation of LH_ox/dyn_ inputs.

Orexin neurons also release glutamate (Rosin et al., 2003). Optogenetic stimulation of LH inputs is likely to induce the concurrent release of other neurotransmitters expressed in orexin neurons. We were unable to evoke AMPAR EPSCs from optical stimulation of LH_ox/dyn_ terminals in the VTA (Extended data 2-1), even though optical stimulation of LH_ox/dyn_ inputs to the VTA can potentiate electrically evoked NMDARs (Thomas et al., 2022). To eliminate potential effects of fast transmission on the activity of VTA^DA^ neurons, evoked firing of VTA^DA^ neurons (n = 12 cells from 9 mice) was recorded in the presence of synaptic blockers including AP5 (50 μM), DNQX (10 μM), and picrotoxin (100 μM), antagonists of NMDA receptors, AMPA receptors, and GABA_A_ receptors, respectively (Fig. 2A,B). As before, we observed that VTA^DA^ neurons were either activated (41% of neurons, RM one way ANOVA: F(4,16)=3.58, p=0.029; Fig. 2B,C) or inhibited (41% of neurons, RM one way ANOVA: F(4,16)=3.74, p=0.025; Fig. 2B,C) by optical stimulation of LH inputs, suggesting that these changes in VTA^DA^ neuron firing are mediated by postsynaptic peptidergic modulation. A subset for neurons were unchanged by optical stimulation (16%, Fig. 2B,C). Increased firing from 30 Hz, 30 s stimulation was significantly different from baseline (p=0.0093) (baseline: 3.6 ± 0.4 APs; 30Hz,30s: 6.6 ± 0.8 APs; SB34867: 5.2 ± 0.9 APs; n/N = 5/3; Fig. 2D), and this change was inhibited by the Ox1R antagonist, SB334867 (1 µM) (p=0.13,RM one-way ANOVA: F(1.516,6.063) = 14.7, p = 0.0059; Fig. 2 D,E). Furthermore, decreased firing from 30 Hz, 30 s stimulation was different from baseline (p=0.037) (baseline: 3.8 ± 0.4 APs; 30 Hz,30 s: 1.8 ± 0.4 APs; NorBNI: 4.2 ± 0.5 APs; n/N = 5/4), and was blocked by the KOR antagonist, NorBNI (1 µM) (p=0.82, RM one-way ANOVA: F(1.764,7.054) = 7.0, p = 0.023; Fig. 2F,G). Because 30 Hz, 30 s optical stimulation of LH_ox/dyn_ inputs to the VTA produced the largest response in either direction, we used this stimulation in the subsequent experiments. Furthermore, all subsequent experiments were conducted in the presence of synaptic blockers.

**Figure 2.**
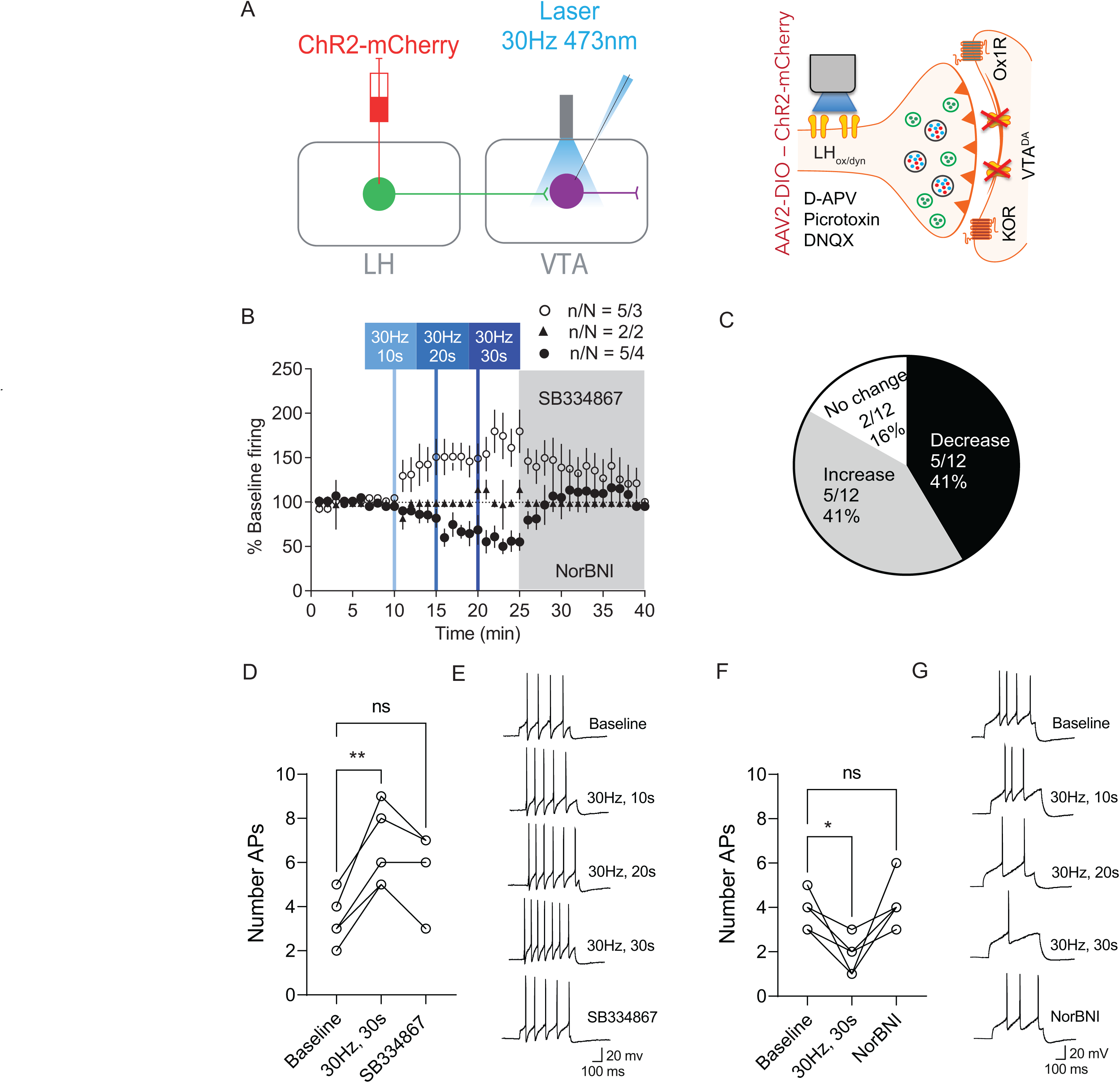
Synaptic inhibitors did not block LH _ox/dyn_ photostimulation induced changes in firing of VTA ^DA^ neurons. **A)** Schematic describing the strategy for optical stimulation of LH inputs on VTA^DA^ neuronal firing in the presence of synaptic blockers including DNQX, AP5, and picrotoxin. Optical stimulation of LH inputs on VTA^DA^ did not induce AMPA oEPSCs – see Figure ED 2-1 for more details. **B)** Time course of evoked firing before and after 30 Hz photoactivation of LH inputs in VTA^DA^ neurons. **C)** Distribution of responses of VTA^DA^ neurons to LH photostimulation. **D)** Number of action potentials during baseline, after 30 Hz, 30 s optical stimulation, and after SB334867 (1 μM) application in VTA^DA^ neurons that increased their response to LH_ox/dyn_ optical stimulation. **E)** Sample traces of evoked action potentials that increased frequency before and after optical stimulation of LH_ox/dyn_ inputs. **F)** Number of action potentials during baseline, after 30 Hz, 30 s optical stimulation, and after NorBNI (1 μM) application in VTA^DA^ neurons that decreased their response to LH optical stimulation. **G)** Sample traces of evoked action potentials that decreased frequency before and after optical stimulation of LH_ox/dyn_ inputs.

### Altered firing of VTA^DA^ neurons upon photoactivation is due to activation of Ox1R and KOR

We next wanted to confirm that optical stimulation of LH_ox/dyn_ inputs to the VTA was releasing orexin or dynorphin leading to activation of their receptors, Ox1R or KOR, respectively expressed in the VTA. In the presence of SB334867 (1 µM), LH optical stimulation decreased VTA^DA^ firing in all neurons (baseline: 100 ± 1%; 30Hz,30s (percent of baseline): 64 ± 5%; NorBNI: 96 ± 6%; n/N=7/4, Fig. 3A). In SB334867, action potentials decreased from baseline (3.7± 0.4) to 2.1 ± 0.3 after 30Hz, 30s stimulation of LH_ox/dyn_ inputs. This effect was washed off by NorBNI (3.6 ± 0.4 APs) (RM one-way ANOVA: F(1.54, 9.14) = 23.54, P = 0.0004), Dunnett’s: Baseline vs. 30Hz, 30s, p=0.0015; Baseline vs. NorBNI, p = 0.96, Fig. 3B,C). We next recorded VTA^DA^ neurons in the presence of NorBNI (1 µM), an antagonist which activates signaling pathways that induce a long lasting suppression of receptor activity and thus is not washed out (Bruchas et al., 2007). LH optical stimulation increased VTA^DA^ firing in five neurons (baseline: 99 ± 0.6%; 30Hz,30s: 126 ± 8%; SB334867: 99.4 ± 0.6%; n/N=5/3) but had no effect in 2 neurons (Fig. 3D). In NorBNI, action potentials increased from baseline (3.9± 0.3) to 4.7 ± 0.5 after 30Hz, 30s stimulation of LH_ox/dyn_ inputs. This effect was washed off by SB334867 (3.9 ± 0.3 APs) (RM one-way ANOVA: F(2, 12) = 10.8, P= 0.0021), Dunnett’s: Baseline vs. 30Hz, 30s, p=0.0032; Baseline vs. SB334867, P> 0.999, Fig. 3E,F). We next recorded VTA^DA^ neurons in the presence of both SB334867 and NorBNI. There was no change in evoked firing of VTA^DA^ neurons after LH optical stimulation (baseline: 103 ± 2%; 30 Hz, 30 s: 105 ± 5%; n=6/5, Fig. 3G). In SB334867 and NorBNI, there was no difference in action potential number between the baseline, (3.2± 0.2), after 30Hz, 30s stimulation of LH_ox/dyn_ inputs (3.3 ± 0.2), and wash (3.2 ± 0.3) (RM one-way ANOVA: F(1.24, 6.34) = 0.55, P = 0.53, Fig. 3H,I). This provides evidence that 1) LH_ox/dyn_ stimulation in the VTA leads to orexin and dynorphin release in the VTA and that 2) increased VTA^DA^ firing after LH optical stimulation is mediated by Ox1R signaling, whereas decreased VTA^DA^ firing is mediated by KOR signaling. Importantly, these effects on firing were found to be independent of synaptic glutamate or GABA release.

**Figure 3.**
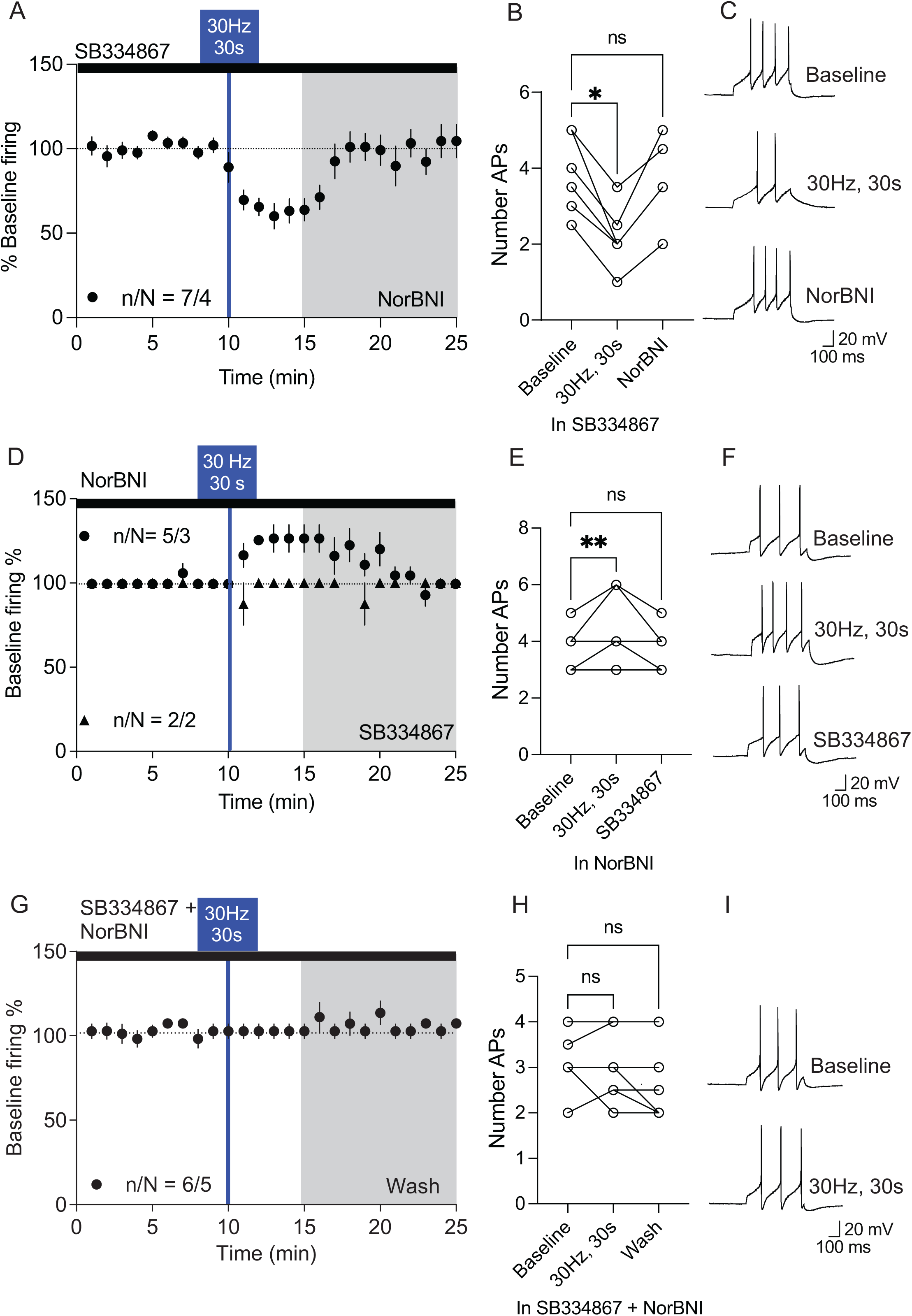
Altered firing of VTA ^DA^ neurons upon photoactivation of LH _ox/dyn_ inputs is due to activation of Ox1Rs and KORs. **A)** All VTA^DA^ neurons decreased firing after photoactivation of LH inputs to the VTA in the presence of synaptic blockers and the Ox1R antagonist, SB334867 (1 μM). **B)** Bar graph of evoked action potentials in the presence of SB334867 and synaptic blockers before and after optical stimulation of LH_ox/dyn_ inputs, and after wash out with NorBNI (1 μM). **C)** Sample traces of evoked action potentials in the presence of SB334867 and synaptic blockers during baseline, after optical stimulation, and after NorBNI treatment. **D)** Time course of evoked firing before and after photoactivation of LH_ox/dyn_ inputs in VTA^DA^ neurons preblocked with NorBNI (1 µM). **E)** LH photoactivation increased the firing of VTA^DA^ neurons preblocked with NorBNI. **F)** Sample traces of evoked action potentials before and after optical stimulation. **G)** Time course of evoked firing before and after photoactivation of LH_ox/dyn_ inputs in VTA^DA^ neurons preblocked with SB334867 and NorBNI (1 µM). **H)** LH photoactivation had no effect on the firing of VTA^DA^ neurons preblocked with SB334867 and NorBNI. **I)** Sample traces of evoked action potentials before and after optical stimulation.

### Temporal characteristics of endogenous LH_ox/dyn_ corelease modulation of DA neuronal activity

Next, we investigated the temporal characteristics of LH -mediated modulation of VTA^DA^ neuronal activity. Following LH optical stimulation, VTA^DA^ neurons that increased or decreased firing were identified and analyzed separately. Increased firing of VTA^DA^ neurons induced by optical stimulation of LH_ox/dyn_ inputs peaked 4 min after stimulation and returned to baseline levels within 25 min (baseline: 99 ± 0.5%; 30Hz,30s: 148 ± 11%; return to baseline, 114 ± 9%; n/N=8/6). We then tested if a second optical stimulation could evoke peptide release. This subsequent stimulation again led to an increase in firing of VTA^DA^ neurons with a peak 4 min after stimulation (2^nd^ optostim, 139 ± 10%). Both first and second peaks after optical stimulation were significantly different to their respective baselines (RM two-way ANOVA: F(1, 7) = 10.17, p = 0.015, Tukey’s multiple comparisons tests: Baseline vs. 30 Hz, 30 s: p=0.0003; 2^nd^ Baseline vs. 2^nd^ optostim: p=0.003; Fig. 4A-C). Conversely, decreased VTA^DA^ firing induced by LH_ox/dyn_ optical stimulation exhibited a peak in response 8 min after stimulation and prolonged inhibition that did not return to baseline after 40 min (baseline: 100 ± 0.8%; 30Hz,30s: 57 ± 8%; n/N=8/5) (Wilcoxon matched-pairs signed rank test: baseline vs optical stimulation, p=0.0156; Fig. 4D-F). When we administered a second optical stimulation 15 min after the first stimulation in VTA^DA^ neurons that decreased firing, we observed a small but significant further decrease in response compared to the second baseline (Fig. 4G-I, RM two-way ANOVA: F(1, 9) = 15.00, p = 0.0038, Tukey’s multiple comparisons tests: Baseline vs. 30 Hz, 30 s: p=0.04; Baseline 2 vs. optostim 2: p = 0.047). In summary, our data reveals a significant difference in the duration of excitatory and inhibitory responses to LH optical stimulation in the VTA, such that inhibition of VTA^DA^ neurons persists, whereas activation of VTA^DA^ neurons is more transient.

**Figure 4.**
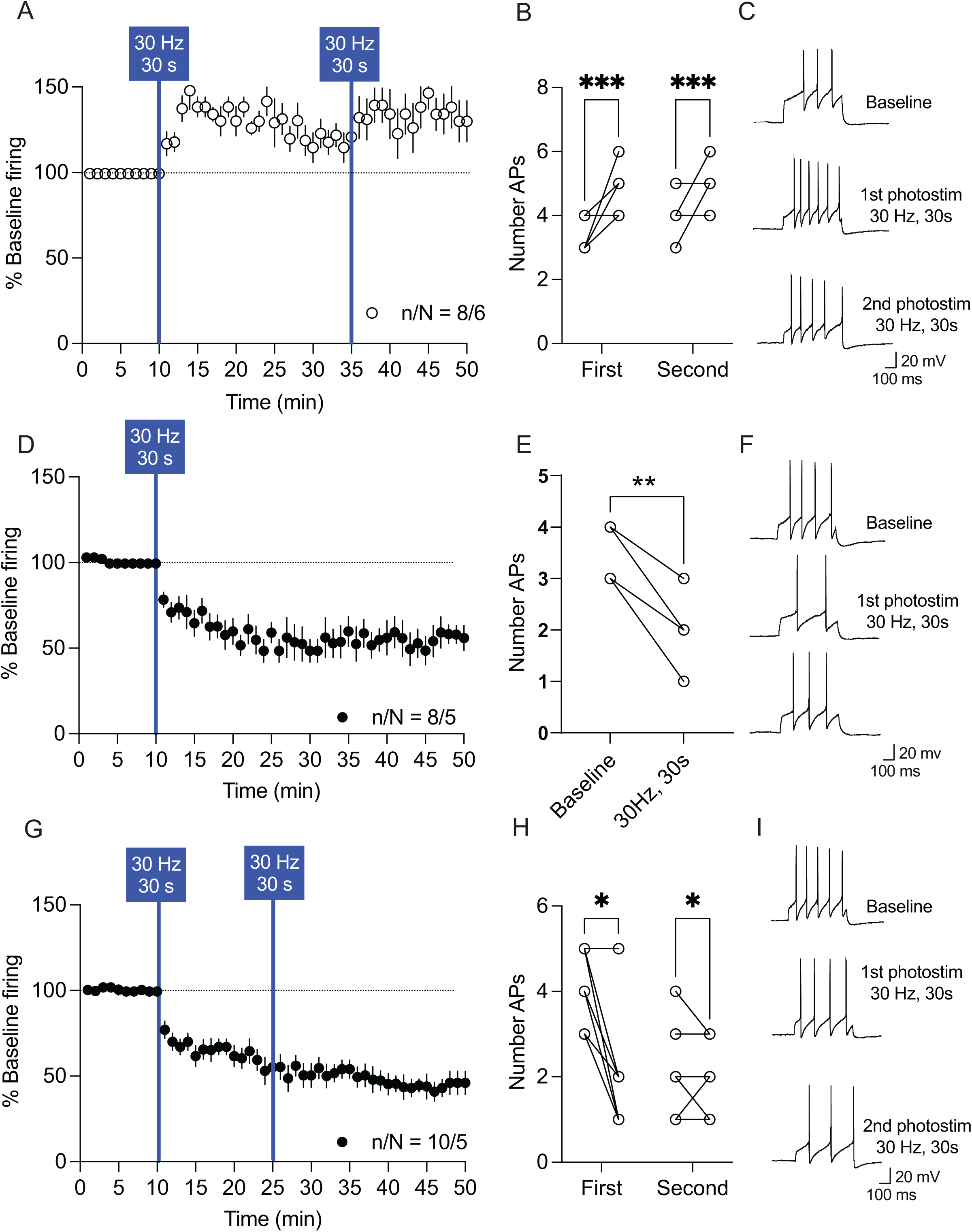
Temporal dynamics of VTA^DA^ neuronal responses to 30 Hz LH_ox/dyn_ optical stimulation. **A)** Time course of evoked firing before and after photoactivation of LH inputs in VTA^DA^ neurons that increased firing. A second photoactivation of LH produced a repeated increase in firing of VTA^DA^ neurons. **B)** Bar graph of evoked action potentials before and after the first and second optical stimulations of LH inputs to VTA^DA^ neurons. **C)** Sample traces of evoked action potentials during baseline, after first and second optical stimulations. **D)** Time course of evoked firing before and after photoactivation of LH inputs in VTA^DA^ neurons that decreased firing. **E)** Bar graph of evoked action potentials before and after optical stimulation of LH inputs to VTA^DA^ neurons that decreased firing. **F)** Sample traces of evoked action potentials during baseline and after optical stimulation. **G)** Time course of evoked firing before and after 2 photoactivations of LH inputs in VTA^DA^ neurons that decreased firing. **H)** Bar graph of evoked action potentials before and after the first and second optical stimulations of LH inputs to VTA^DA^ neurons that decreased firing. **F)** Sample traces of evoked action potentials during baseline, after the first and second optical stimulations.

### Photoactivation of LH _ox/dyn_ terminals VTA DA has diverse effects on evoked firing based on projection target

The VTA^DA^ system is heterogeneous and is increasingly thought about in terms of anatomically and functionally distinct sub-networks (Watabe-Uchida et al., 2012). VTA^DA^ neurons project to different regions on the basis of their localization along the mediolateral axis (Lammel et al., 2008; Beier et al., 2015, 2019). We next tested if the distinct firing responses induced by optical stimulation of LH_ox/dyn_ inputs segregate by dopaminergic projection target. Therefore, in orexin^cre^ mice expressing ChR2 in orexin neurons, we recorded from VTA^DA^ neurons retrogradely labeled from the subregions of the NAc or the BLA using fluorescent beads. We first targeted the NAc by injecting red Lumifluor RetroBeads, a retrograde tracer, in two sub-nuclei: the lateral shell (IAcbSh) (Fig. 5 A-C) and the medial shell (mAcbSh) of orexin^cre^ mice expressing ChR2 in LH neurons (Fig. 6 A-C). We confirmed the retrobead injection sites (Fig. 5C, 6C, 7C), as well as the TH expression in the VTA neuron projecting to the lAcbSh (Fig. 5A), mAcbSh (Fig. 6A), or BLA (Fig. 7A). We also compared electrophysiological characteristics of VTA^DA^ neurons with known projections (Supplemental Fig. 5-1). As reported previously (Lammel et al., 2011; Baimel et al., 2017), VTA^DA^ neurons that project to the lAcbSh have larger hyperpolarization-activated current (Ih) than those projecting to the mAcbSh or the BLA (Kruskall-Wallis test, p = 0.0079) with significant differences between VTA^DA^ neurons that project to the lAcbSh and mAcbSh (p = 0.016) or BLA (P = 0.024) (Dunn’s multiple comparison test; Fig. 5-1A). There was also a significant difference between groups on capacitance (Kruskall-Wallis test, p = 0.0009), with VTA^DA^ neurons that project to the lAcbSh having a larger capacitance, reflecting larger cell size, than those projecting to the mAcbSh (p = 0.0012) or the BLA (p = 0.0135, Dunn’s multiple comparison test; Fig. 5-1B). Input resistance was also different between groups (Kruskall-Wallis test, p = 0.0175), with a significant difference between VTA^DA^ neurons that project to the lAcbSh and the BLA (p = 0.02, Dunn’s multiple comparison’s test; Fig. 5-1C). Taken together VTA^DA^ neurons projecting to the lAcbSh have larger Ih current and capacitance and smaller input resistance than those projecting to the mAcbSh or the BLA.

**Figure 5.**
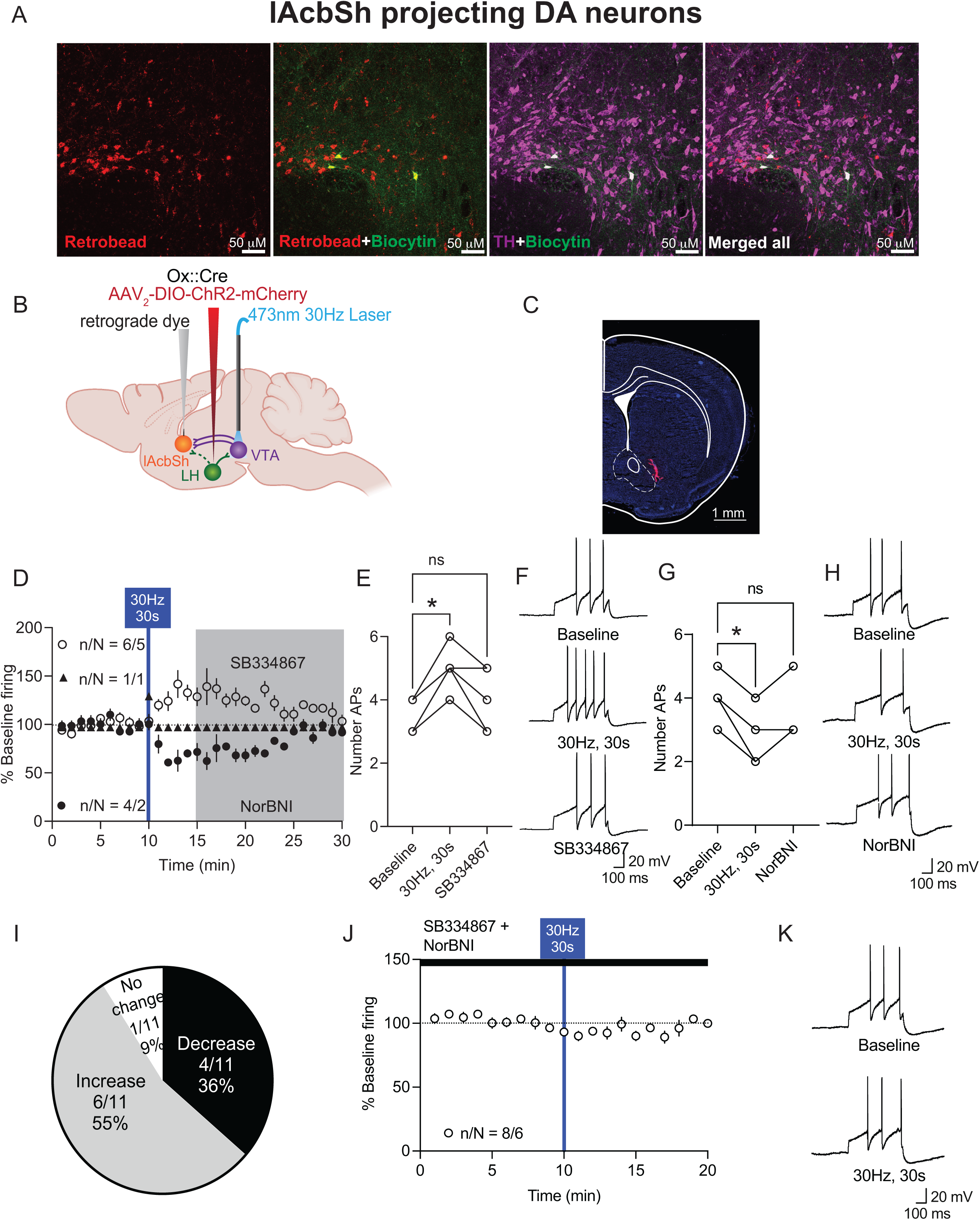
Photoactivation of LH _ox/dyn_ inputs preferentially activates most lAcbSh-projecting VTA ^DA^ neurons. **A)** Images of VTA sections with TH positive neurons (purple), biocytin (green), and retrograde labeled (red) neurons from the lAcbSh. Scale bar, 50 µm. **B)** Schematic of viral strategy **C)** Example of RetroBead injection site in the lAcbSh of orexin^cre^ mice. Scale bar, 1 mm. **D)** Time course of evoked firing before and after photoactivation of LH inputs in lAcbSh projecting VTA^DA^ neurons. **E)** Evoked action potentials before and after optical stimulation of LH to lAcbSh-projecting VTA^DA^ neurons, and in the presence of SB334867 for VTA^DA^ neurons that increased their firing to LH stimulation. LAcbSh-projecting VTA^DA^ neurons have different electrophysiological properties that BLA- or mAcbSh-projectiving VTA^DA^ neurons. See Figure ED 5-1 for more details. **F)** Sample traces of evoked action potentials before and after optical stimulation for VTA^DA^ neurons that increased firing to photostimulation of LH inputs. **G)** Evoked action potentials before and after optical stimulation of LH to lAcbSh-projecting VTA^DA^ neurons, and in the presence of NorBNI for VTA^DA^ neurons that decreased their firing to LH stimulation. **H)** Sample traces of evoked action potentials before and after optical stimulation for VTA^DA^ neurons that decreased firing to photostimulation of LH_ox/dyn_ inputs. **I)** Distribution of responses of lAcbSh-projecting VTA^DA^ neurons to LH photostimulation. **J)** Time course of evoked firing of lAcbSh-projecting VTA^DA^ neurons preblocked with SB334867 and NorBNI. **K)** Sample traces of evoked action potentials of lAcbSh-projecting VTA^DA^ neurons before and after optical stimulation in the presence of SB334867 and NorBNI.

**Figure 6.**
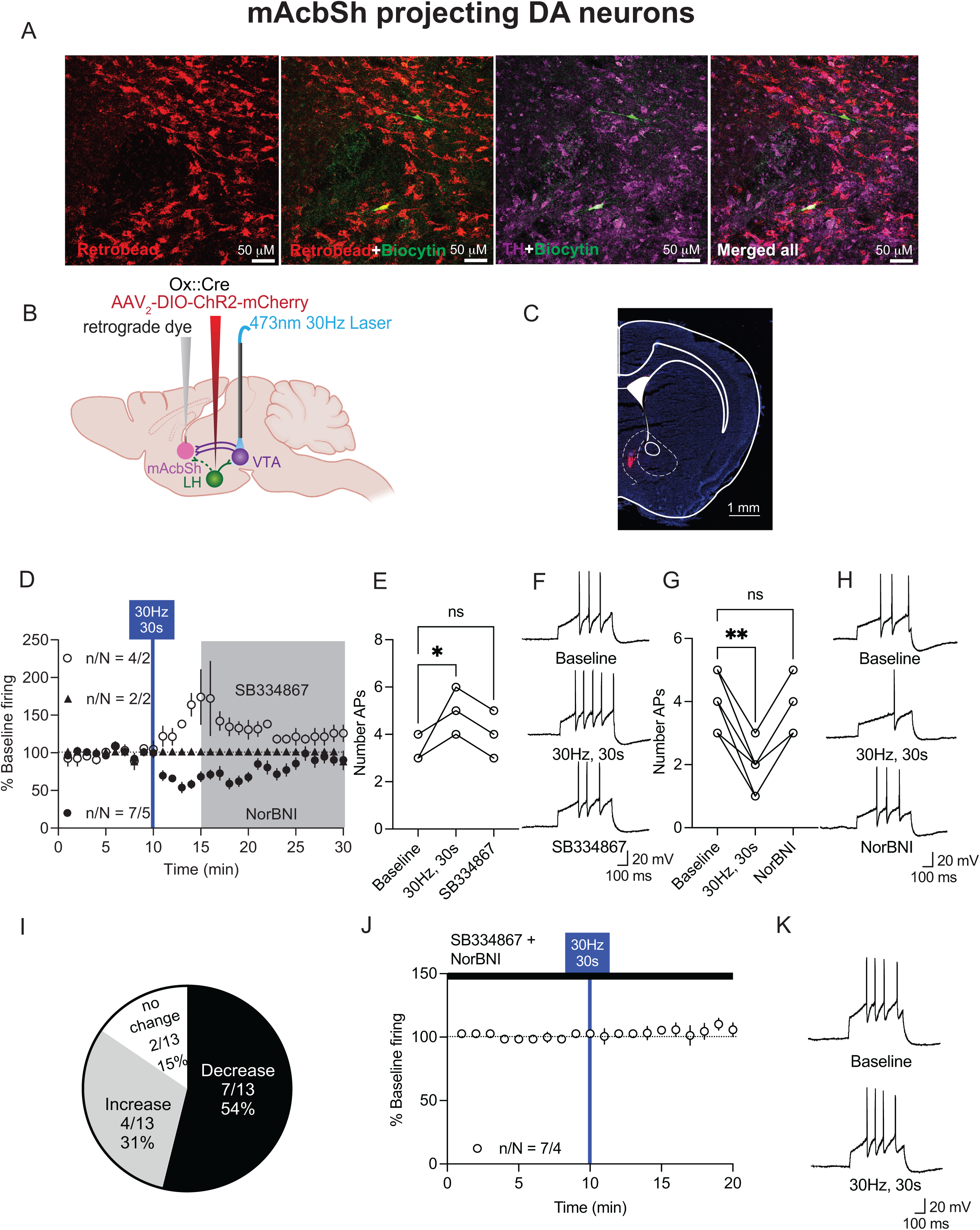
Photoactivation of LH _ox/dyn_ inputs bidirectionally affects the neuronal firing of mAcbSh-projecting VTA^DA^ neurons. **A)** Images of VTA sections with TH positive neurons (purple), biocytin (green), and retrograde labeled (red) neurons from the mAcbSh. Scale bar, 50 µm. **B)** Schematic of viral strategy **C)** Example of RetroBead injection site in the mAcbSh of orexin^cre^ mice. Scale bar, 1 mm. **D)** Time course of evoked firing before and after photoactivation of LH inputs in mAcbSh projecting VTA^DA^ neurons. **E)** Evoked action potentials before and after optical stimulation of LH to mAcbSh-projecting VTA^DA^ neurons, and in the presence of SB334867 for VTA^DA^ neurons that increased their firing to LH stimulation. **F)** Sample traces of evoked action potentials before and after optical stimulation for VTA^DA^ neurons that increased firing to photostimulation of LH_ox/dyn_ inputs. **G)** Evoked action potentials before and after optical stimulation of LH_ox/dyn_ to mAcbSh-projecting VTA^DA^ neurons, and in the presence of NorBNI for VTA^DA^ neurons that decreased their firing to LH_ox/dyn_ stimulation. **H)** Sample traces of evoked action potentials before and after optical stimulation for mAcbSh-projecting VTA^DA^ neurons that decreased firing to photostimulation of LH inputs. **I)** Distribution of responses of mAcbSh-projecting VTA^DA^ neurons to LH photostimulation. **J)** Time course of evoked firing of mAcbSh-projecting VTA^DA^ neurons preblocked with SB334867 and NorBNI. **K)** Sample traces of evoked action potentials of mAcbSh-projecting VTA^DA^ neurons before and after optical stimulation in the presence of SB334867 and NorBNI.

**Figure 7.**
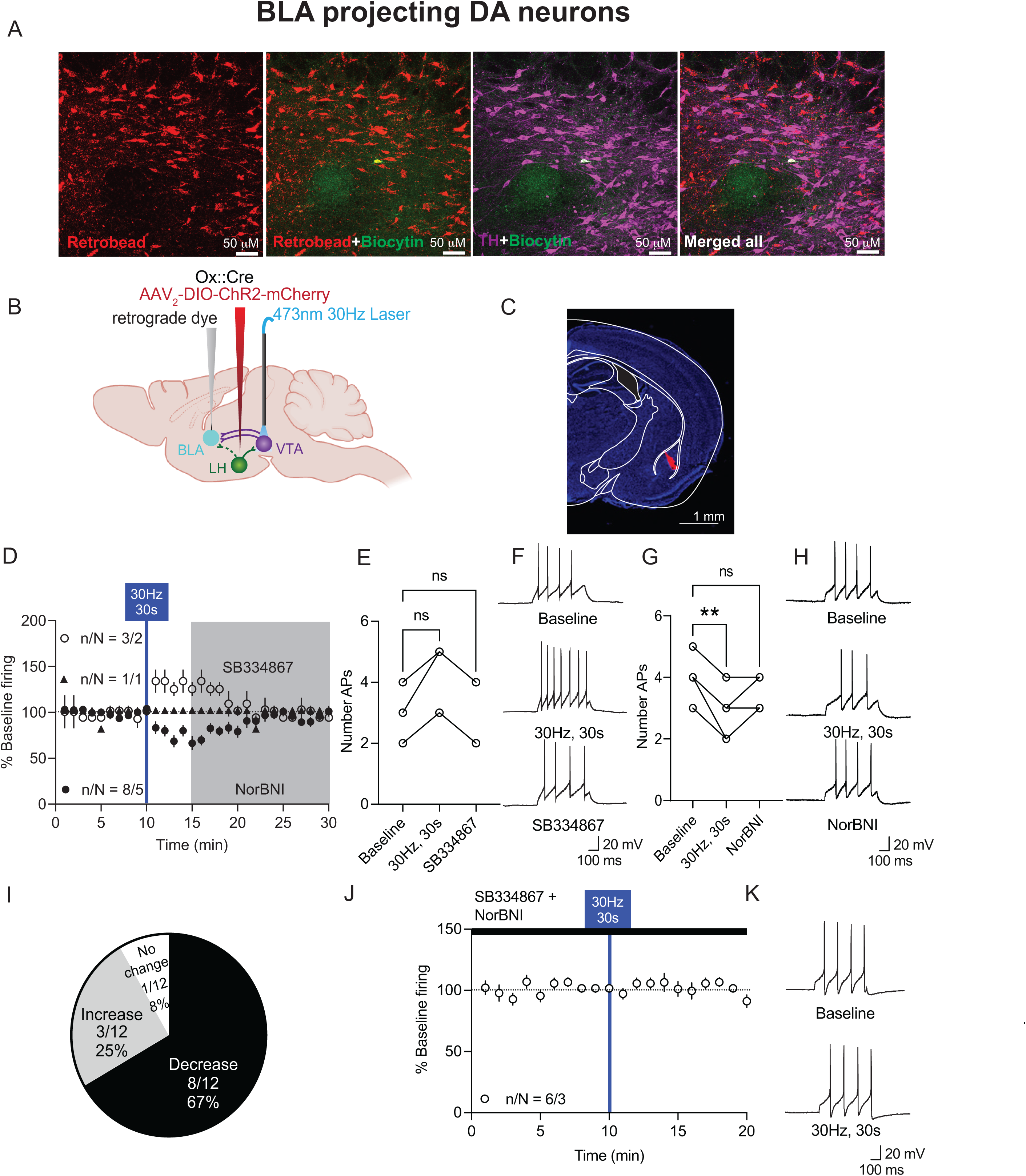
Photoactivation of LH_ox/dyn_ inputs to BLA-projecting VTA^DA^ primarily decreases firing. **A)** Images of VTA sections with TH positive neurons (purple), biocytin (green), and retrograde labeled (red) neurons from the BLA. Scale bar, 50 µm. **B)** Schematic of viral strategy **C)** Example of RetroBead injection site in the BLA of orexin^cre^ mice. Scale bar, 1 mm. **D)** Time course of evoked firing before and after photoactivation of LH inputs in BLA-projecting VTA^DA^ neurons. **E)** Evoked action potentials before and after optical stimulation of LH to BLA-projecting VTA^DA^ neurons, and in the presence of SB334867 for VTA^DA^ neurons that increased their firing to LH stimulation. **F)** Sample traces of evoked action potentials before and after optical stimulation for VTA^DA^ neurons that increased firing to photostimulation of LH_ox/dyn_ inputs. **G)** Evoked action potentials before and after optical stimulation of LH to BLA-projecting VTA^DA^ neurons, and in the presence of NorBNI for VTA^DA^ neurons that decreased their firing to LH_ox/dyn_ stimulation. **H)** Sample traces of evoked action potentials before and after optical stimulation for BLA-projecting VTA^DA^ neurons that decreased firing to photostimulation of LH inputs. **I)** Distribution of responses of BLA-projecting VTA^DA^ neurons to LH photostimulation. **J)** Time course of evoked firing of BLA-projecting VTA^DA^ neurons preblocked with SB334867 and NorBNI. **K)** Sample traces of evoked action potentials of BLA-projecting VTA^DA^ neurons before and after optical stimulation in the presence of SB334867 and NorBNI.

We next characterized the evoked firing responses of VTA^DA^ neurons to optical stimulation of LH inputs. We found that 55% of VTA^DA^ neurons projecting to the lAcbSh increased firing to optical stimulation of LH_ox/dyn_ inputs, whereas 36% decreased firing (Fig. 5D-I). Of 11 DA-lAcbSh neurons, 6 neurons increased firing (baseline: 3.5 ± 0.2 APs; 30Hz, 30s: 4.8 ± 0.3 APs; SB334867: 3.8 ± 0.4 APs; n/N=6/5 mice, RM one way ANOVA: F(2,15) = 4.72, P = 0.0256, Dunnett’s: baseline vs 30Hz, 30s p = 0.018, baseline vs SB334867: p = 0.69; Fig. 5E,F). Increases in firing were blocked by SB33864 (Fig. 5E), suggesting increases in firing were mediated by Ox1R signaling. Four of 11 neurons decreased firing in response to optical stimulation of LH_ox/dyn_ inputs (baseline: 4 ± 0.4 APs; 30 Hz, 30 s: 2.7 ± 0.5 APs; NorBNI: 3.5 ± 0.5 APs; n/N=4/2 mice, RM one way ANOVA: F(1.92,5.78) = 11.4, P = 0. 01; Dunnett’s: baseline vs 30 Hz, 30 s: p = 0.025, baseline vs NorBNI: p = 0.28; Fig. 5G,H). Decreased firing was blocked by NorBNI (Fig. 5G), suggesting, that this was mediated by KOR signaling. One lAcbSh-projecting VTA^DA^ neuron exhibited no change in firing (Fig 5.D, I). In the presence of Ox1R and KOR antagonists, there was no change in firing of VTA^DA^ neurons after LH optical stimulation (Fig. 5J,K), suggesting changes in firing were mediated by peptide release.

The 30 Hz optical stimulation of LH_ox/dyn_ inputs in the VTA had a similar bidirectional effect on mAcbSh projecting VTA^DA^ neurons. Four out of 13 (31%) VTA^DA^ neurons recorded increased firing in response to LH_ox/dyn_ stimulation (baseline: 3.5 ± 0.3 APs; 30 Hz, 30 s: 5.0 ± 0.4 APs; SB334867: 4.0 ± 0.4 APs; n/N=4/2 mice, RM one way ANOVA: F(1.39,4.19) = 9.0, P = 0.034, Dunnett’s: baseline vs 30Hz, 30s p = 0.05, baseline vs SB334867: p = 0.56; Fig. 6E,F). Additionally, the stimulation led to a decrease in firing activity for 7 out of 13 (54%) recorded neurons (baseline: 4.1 ± 0.3 APs; 30 Hz, 30 s: 2.0 ± 0.3 APs; NorBNI: 3.6 ± 0.3 APs; n/N=7/5 mice, RM one way ANOVA: F(1.35,8.1) = 66.3, P < 0. 0001; Dunnett’s: baseline vs 30 Hz, 30 s p = 0.0003, baseline vs NorBNI: p = 0.28; Fig. 6G,H). The increase in firing was reversed by SB334867 (Fig. 6E,F), whereas the decrease in firing was reversed by NorBNI (Fig. 6G,H). Two of 13 cells had no response (15%; Fig. 6D,I). There was no change in firing after optical stimulation of LH inputs to VTA^DA^ neurons in the presence of both KOR and Ox1R antagonists (Fig. 6J,K), confirming that bidirectional changes in firing are mediated by LH orexin or dynorphin.

We next examined the response of BLA projecting VTA^DA^ neurons to LH optical stimulation. The majority of BLA-projecting VTA^DA^ (8 of 12; 67%) are inhibited by optical stimulation of LH inputs (baseline: 3.9 ± 0.2 APs; 30Hz, 30s: 2.7 ± 0.2 APs; NorBNI: 3.4 ± 0.2 APs; RM one-way ANOVA: F(1.89, 15.06) = 31, P < 0.0001, Dunnett’s: baseline vs. 30Hz, 30s, p=0<0.0001; baseline vs. NorBNI: p = 0.06; Fig. 7D,G,H). This inhibition of firing was blocked by NorBNI (Fig. 7D,G). Optical stimulation increased firing in only 3 of 12 (25%) BLA-projecting VTA^DA^ neurons although this did not reach statistical significance (baseline: 3 ± 0.6 APs; 30Hz, 30s: 4.3 ± 0.7 APs; SB334867: 3.3 ± 0.7; RM one-way ANOVA: F(1.0, 2.0) = 13.0, P = 0.069; Fig. 7D,E,F,I). One of 12 neurons recorded exhibited no change (Fig 7D,I). Finally, there was no change in firing after optical stimulation of LH inputs to BLA-projecting VTA^DA^ neurons in the presence of both KOR and Ox1R antagonists (Fig. 7J,K). Taken together, 30 Hz LH_ox/dyn_ photoactivation inhibited a large proportion of BLA-projecting VTA^DA^ neurons but fewer lAcbSh-projecting or mAcbSh-projecting VTA^DA^ neurons. Overall, these results indicate that the majority of VTA^DA^ neurons activated by LH_ox/dyn_ stimulation project to the lAcbsh, whereas the majority of inhibited neurons project to the BLA or the mAcbSh. These experiments identify how firing of lAcbSh, mAcbSh, or BLA-projecting VTA^DA^ neurons can be tuned by co-release of neuropeptides from the LH in a physiological state, such that LH_ox_ dominates in the VTA^DA^-lAcbSh projection whereas LH_dyn_ dominates in the VTA^DA^-mAcbSh and BLA projections.

## Discussion

Here, we demonstrated that optical stimulation of LH _ox/dyn_ inputs have distinct modulatory effects on the activity of projection-target-defined VTA^DA^ neurons. We showed that 30 Hz optical stimulation of LH produced both orexin and dynorphin neuromodulation of VTA^DA^ firing. LH stimulation-induced increased firing was blocked by the Ox1R antagonist, whereas LH_ox/dyn_ stimulation induced decrease firing was blocked by the KOR antagonist. When both peptide receptors were blocked, no change in firing occurred after LH_ox/dyn_ stimulation, suggesting that changes in firing are mediated by peptide release into the VTA. Furthermore, optical stimulation of LH_ox/dyn_ inputs increased firing in lAcbSh-projecting VTA^DA^ neurons, but mostly inhibited firing in the mAcbSh and BLA projections. These results suggest that corelease of orexin and dynorphin may balance ensembles of VTA^DA^ within each projection to influence dopamine output.

Several whole animal behavioral studies have demonstrated optogenetic stimulation of behavioral responses consistent with peptide release (Adamantidis et al., 2007; Atasoy et al., 2012; Jego et al., 2013; Kempadoo et al., 2013). However, the temporal dynamics of the release and the postsynaptic neurons upon which this neuromodulatory action occurs is not clear. This can be addressed in brain slice preparations where optical stimulation frequency and duration can be controlled and responses to fast acting amino-acid neurotransmitters or co-released peptides can occur. The LH_ox/dyn_ input to the VTA presents a unique preparation where only a small proportion of terminals synapse onto VTA neurons, but there is a high density of peptide-containing dense core vesicles within the VTA, allowing for neuromodulatory action from peptide release (Balcita-Pedicino and Sesack, 2007). While single brief (2-5 ms) light pulses are sufficient to evoke neurotransmitters from synaptic vesicles as voltage dependent calcium channels are highly coupled to synaptic vesicles in the active zone, these protocols may be insufficient to release neuropeptides (Bruns and Jahn, 1995; Leenders et al., 1999; Südhof, 2012). The release of neuropeptides typically requires a higher frequency and longer duration of depolarization to allow for sufficient calcium-dependent mobilization of dense core vesicles to the plasma membrane (Weisskopf et al., 1993; Leenders et al., 1999; Muschol and Salzberg, 2000). This may be related to the location of dense core vesicles away from the active zone (Bruns and Jahn, 1995; van den Pol, 2012). We found that optical stimulation at 30 Hz with a 5 ms pulse width for either 10, 20, or 30 seconds could alter the firing rate of VTA^DA^ neurons. Importantly, this occurred in the absence of amino acid mediated synaptic transmission and was blocked by antagonists for Ox1R and KOR, suggesting neuropeptide release into the VTA. Consistent with this, we found that optogenetically stimulating glutamatergic LH orexin neurons did not produce AMPA excitatory postsynaptic currents, but potentiated electrically evoked NMDA currents, suggesting a neuromodulatory role (Thomas et al., 2021). Thus, we demonstrate here a bidirectional neuromodulatory effect of LH orexin and dynorphin on VTA^DA^ firing.

Orexin neurons have intrinsic features that promote long-lasting firing activity (Burt et al., 2011). Orexin neurons are in an intrinsically depolarized state largely due to the constitutively active cation current by transient receptor potential C channels (Cvetkovic-Lopes et al., 2010). This depolarized state maintains neurons near their firing threshold leading to sustained spontaneous firing. In vivo recordings suggest that orexin neurons exhibit slow (<10 Hz) tonic discharges during wakefulness (Takahashi et al., 2008; Hassani et al., 2009). These mechanisms are likely important for the physiological functions of orexin neurons, which require prolonged output to maintain a wakeful state. However, orexin neurons can also follow faster frequencies likely important for the release of neuropeptides from dense core vesicles. For example, action potentials of LH_ox/dyn_ neurons can efficiently follow optogenetic stimulation at 30 Hz (Thomas et al., 2022) up to 50 Hz frequencies (Adamantidis et al., 2007). Several local mechanisms might contribute to this elevated firing activity. Orexin can be somatodentrically released which activates Ox2Rs to open non-selective cation channels which can depolarize orexin neurons or increase presynaptic glutamate release (Li et al., 2002). Furthermore, orexin neurons receive numerous glutamatergic inputs which outnumber inhibitory synapses that may also contribute to sustained depolarized states required for neuropeptide release (Horvath and Gao, 2005). Finally, orexin neurons use astrocyte-derived lactate as an energy substrate to maintain spontaneous firing and the excitatory action of glutamatergic transmission (Parsons and Hirasawa, 2010). Circumstances in which these sustained frequencies, and presumed orexin release, may occur in response to arousing, motivationally relevant situations. Recordings of presumed orexin neurons in freely moving rats found that orexin neurons have tonic activity during active waking, grooming, and eating, but have rapid firing during periods of adaptive behaviours, such as exploration, play, and predation (Mileykovskiy et al., 2005; Wu et al., 2011). Highly arousing events such as stress and reward-seeking, may also sufficiently increase sustained firing to promote neuropeptide release, which may be required to engage monoamine neuromodulatory systems to either escape or take advantage of the opportunity (Giardino and de Lecea, 2014). Thus, orexin release may occur in response to salient, motivationally relevant events to coordinate adaptive behaviours (reviewed in (Mahler et al., 2014).

Application of the Ox1R antagonist, SB334867, inhibited the excitatory effects of LH_ox/dyn_ stimulation. Notably, orexin can interact with both Ox1R and Ox2R that are expressed within the VTA (Marcus et al., 2001; Narita et al., 2006). The affinity of SB334867 is 40 nM at Ox1R which is 50 fold selective over Ox2R (Smart et al., 2001); however a bath application of 1 μM likely has inhibitory action at both Ox1R and Ox2R. Ox1R and Ox2R are typically expressed postsynaptically on VTA^DA^ and some GABAergic neurons (Fadel and Deutch, 2002; Balcita-Pedicino and Sesack, 2007), although some reports have demonstrated a presynaptic action of orexin A in the VTA (Borgland et al., 2009). Although few Gq coupled receptors are expressed presynaptically, Ox1 an Ox2 receptors are known to be promiscuous in their G protein alpha subunit coupling (Kukkonen and Leonard, 2014). KORs are also expressed on both somatodendrites of VTA^DA^ neurons but mediate their postsynaptic effects via opening a G-protein coupled inwardly rectifying potassium conductance via activation of Gi/o-coupled KORs. KORs can also be expressed at terminals within the VTA, such that dynorphin can suppress both excitatory and inhibitory synaptic transmission onto VTA^DA^ neurons (Margolis et al., 2005; Ford et al., 2006). Importantly, in these experiments, photostimulation of LH_ox/dyn_ activated or inhibited firing in the presence of synaptic blockers, thus negating possible presynaptic or indirect activation. Taken together, neuromodulatory action of LH photostimulation of VTA^DA^ neurons was mediated by postsynaptic somatodendritic Ox1R or KOR activation.

The time courses for the LH neuromodulatory effect on VTA^DA^ neurons were different for either peptide. LH_ox/dyn_ optical stimulation produced a sustained decrease in firing. Because this effect could be washed off with NorBNI, it suggests that dynorphin lingers in the slice possibly due to decreased metabolic peptidases for dynorphin that maybe washed out with continuous superfusion of the slices. Exogenous application of dynorphin (5 min) to VTA slices in the presence of peptidase inhibitors, captopril and beestatin, also produced a lasting depression of VTA^DA^ firing that did not return to baseline 15 min after washout (Baimel et al., 2017). This is in contrast to a small molecule KOR agonist, U69593, that has transient effects in the VTA (Margolis et al., 2003), suggesting that the lasting effect is unlikely to be due to receptor kinetics. Notably, dynorphin application in the absence of peptidase inhibitors to orexin neurons also produced a transient inhibition of firing (Li and van den Pol, 2006). Taken together, we speculate that the lasting effects of dynorphin release on VTA^DA^ firing are due to poor washout and decreased metabolism of the peptide in the slice preparation. In contrast, the effects of orexin were shorter lasting, with a significant decay in response 20 min after application, likely due to faster degradation or washout of orexin. Additionally, the differential effects of dynorphin and orexin on VTA^DA^ firing can be attributed to their distinct signaling mechanisms (Bruchas and Chavkin, 2010; Kukkonen and Leonard, 2014). Exogenous application of orexin (5 min) also has a longer lasting increase of VTA^DA^ firing, elevated 15 min after application (Baimel et al., 2017). Notably, we were able to evoke presumed orexin release for a second time in VTA slices. This suggests that there are sufficient dense core vesicles to mobilize for repeated release, which may have implications for how LH_ox/dyn_ neurons can signal in regions with high density of vesicles. The smaller inhibition with a second stimulation of LH_ox/dyn_ is likely because of a ceiling effect of KOR activation.

Previous work has demonstrated that exogenous application of orexin and dynorphin modulate non-overlapping DAergic circuits originating from the VTA to tune dopaminergic output (Baimel et al., 2017). Specifically, exogenous application of orexin potentiates firing of VTA^DA^ neurons that project to the lAcbSh, but not the BLA, whereas exogenous dynorphin inhibits firing in subpopulations of DA neurons in both mAcbSh and lAcbSh and inhibits most BLA projecting DA neurons (Baimel et al., 2017). While LH orexin neurons provide the only source of orexin to the VTA (Peyron et al., 1998), dynorphin is also expressed in the substantia nigra, prefrontal cortex, ventral and dorsal striatum, the central amygdala, and dorsal raphe (Watson et al., 1982; Weber et al., 1982; Fallon and Leslie, 1986; Healy and Meador-Woodruff, 1994; Abraham et al., 2022). The dynorphin expressing ventral striatal and dorsal raphe neurons project to the VTA and supply a dynorphin modulatory input in addition to that of LH_ox/dyn_ neurons (Shippenberg et al., 2001; Abraham et al., 2022). Given that LH_ox/dyn_ neurons are activated during arousing situation, such as stress or motivationally advantageous opportunities, it is important to determine how neuropeptides from this particular input can tune selective VTA originating circuits. We found that, like exogenous application, LH dynorphin decreased the activity of the majority of BLA – projecting VTA^DA^ neurons (Baimel et al., 2017). BLA DA can shape attention related learning signals and is involved in encoding identity specific cue memories (Esber et al., 2012; Sias et al., 2024). Thus, a suppression of this input during periods when LH_ox/dyn_ neurons are activated may reduce formation of these memories, although further research is required to test this hypothesis.

We found that LH activation of NAc projecting VTA^DA^ neurons had differential effects dependent on NAc shell subregion. Whereas the majority of VTA^DA^ neurons that projected to the lAcbSh were activated by LH stimulation, VTA^DA^ neurons projecting to the mAcbSh were primarily inhibited. Although both projections had a large proportion of neurons that also responded to dynorphin, consistent with studies using exogenously applied dynorphin (Ford et al., 2006; Baimel et al., 2017). This differentiation may arise, to some extent, from differences in the expression of Ox1R or KOR on VTA neurons projecting to the mAcbSh vs the lAcbSh. However, conclusive evidence for this hypothesis is yet to be established. Taken together, one reason why orexin and dynorphin may be co-expressed and co-released, is to simultaneously modulate different VTA^DA^ ensembles that can then influence their downstream projections.

VTA^DA^ neurons exhibit heterogeneity in their axonal projections, electrophysiological characteristics, and various molecular features. However, the functional consequences of this diversity on behavior remains poorly elucidated. DA projections from the VTA to the NAc, play an important role in motivated behaviors, reinforcement learning, and reward processing (Hamid et al., 2016; Kim and Kaang, 2022). Recent studies have shown the contribution of VTA projections to the amygdala in encoding state-specific motivational salience (Lutas et al., 2019), regulating approach/avoidance behavior toward threats (Miller et al., 2019), and have established the role of VTA projections in modulating BLA activity during aversive conditioning (Tang et al., 2020). An interesting example of how VTA^DA^ projections to the NAc or the amygdala are influenced in response to nicotine administration revealed that a nicotine injection induces opposing responses in two distinct subpopulations of VTA^DA^ neurons (Nguyen et al., 2021). Specifically, a majority of lateral VTA^DA^ projections to the NAc are activated, while a considerable majority of medial VTA^DA^ neurons with axons projecting to the amygdala are inhibited after systemic nicotine (Nguyen et al., 2021). Nguyen et al. (2021) also demonstrated that both rewarding and anxiogenic effects of nicotine exposure occur simultaneously and are conveyed by distinct subpopulations of VTA^DA^ neurons, such that inhibition of amygdala-projecting VTA^DA^ neurons mediates anxiety-like behaviour and their activation prevents the anxiogeneic effect of nicotine, whilst activation of NAc projecting VTA^DA^ neurons likely mediates the reinforcing effect of nicotine (Nguyen et al., 2021). Future research should examine how the concurrent engagement of two circuits with opposing messages could compete to produce specific functional outcomes and whether an imbalance between the two could lead to brain disorders such as addiction.

In summary, we demonstrate that BLA-projecting VTA^DA^ neurons were predominantly inhibited in response to LH optical stimulation, whereas lAcbSh-projecting and mAcbSh-projecting VTA^DA^ neurons were bidirectionally modulated. Thus, it would be predicted that corelease of LH orexin and dynorphin might balance ensembles of VTA^DA^ neurons within different projections to influence their final output. Given that orexin signaling in the VTA drives motivated reward seeking, we speculate that the effects of orexin and dynorphin corelease in the VTA may act to coordinate dopaminergic output to bias activity toward projection targets like the NAc, which are critical for effort-driven reward seeking (Correa et al., 2002), while dampening activity in other circuits that are less essential in these tasks.

## Acknowledgements

The authors would like to acknowledge the Hotchkiss Brain Institute advanced microscopy facility. This research was performed at the University of Calgary which is located on the unceded traditional territories of the people of the Treaty 7 region in Southern Alberta, which includes the Blackfoot Confederacy (including the Siksika, Piikuni, Kainai First Nations), the Tsuut’ina, and the Stoney Nakoda (including the Chiniki, Bearspaw, and Goodstoney First Nations). The City of Calgary is also home to Metis Nation of Alberta, Region III.

## Author Contributions

AM performed surgeries and electrophysiological experiments. MQ performed immunohistochemistry and imaging experiments. AM, MQ, and SLB analyzed the data. AM and SLB wrote first drafts of the manuscript. AM, MQ, and SLB revised the manuscript.

---

**Figure ED 2-1.**
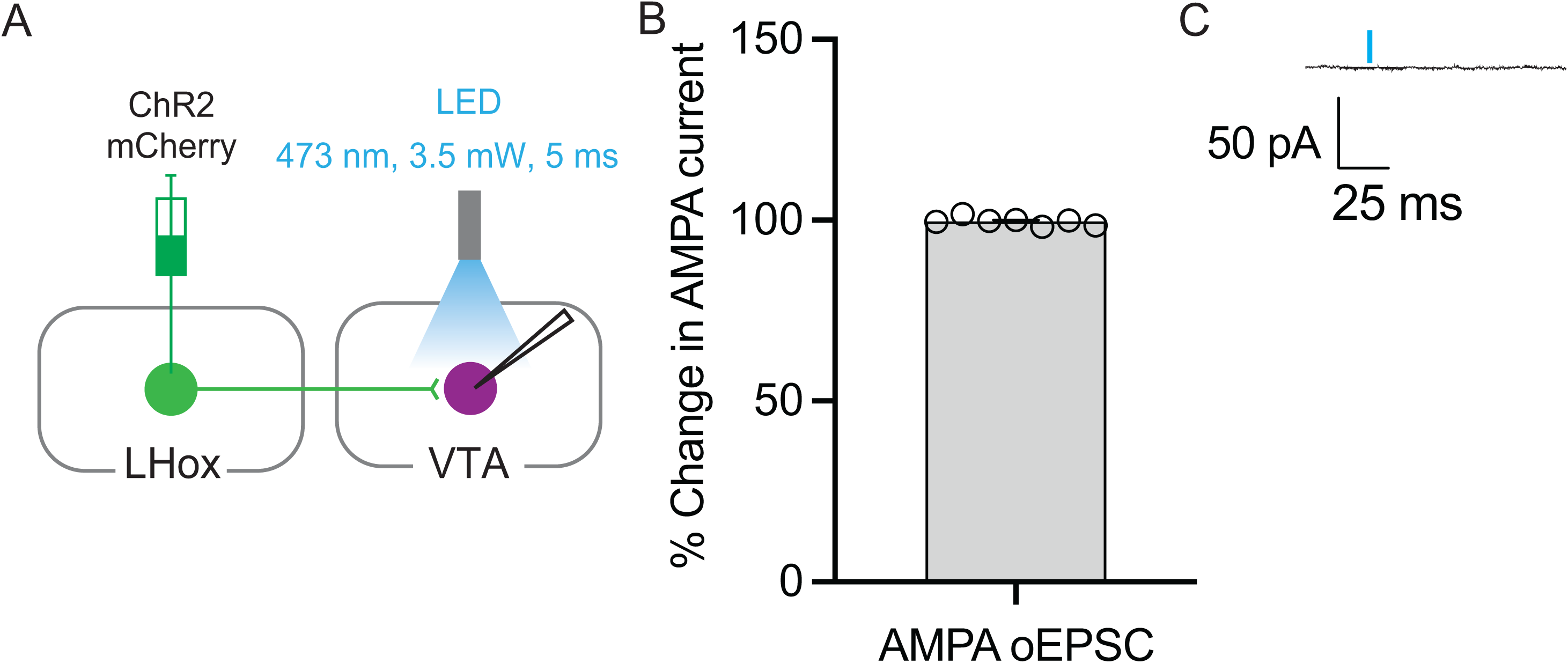
Optical stimulation of LH_ox/dyn_ inputs does not produce AMPA EPSCs in the VTA. **A)** Diagram of parameters used to optically stimulate AMPA EPSCs recorded at -70 mV in the presence of picrotoxin in the VTA. **B)** Percent change in response post optical stimulation compared to pre-stimulation baseline. **C)** Example sweep from a neuron recorded before and after optical stimulation.

**Figure ED 5-1.**
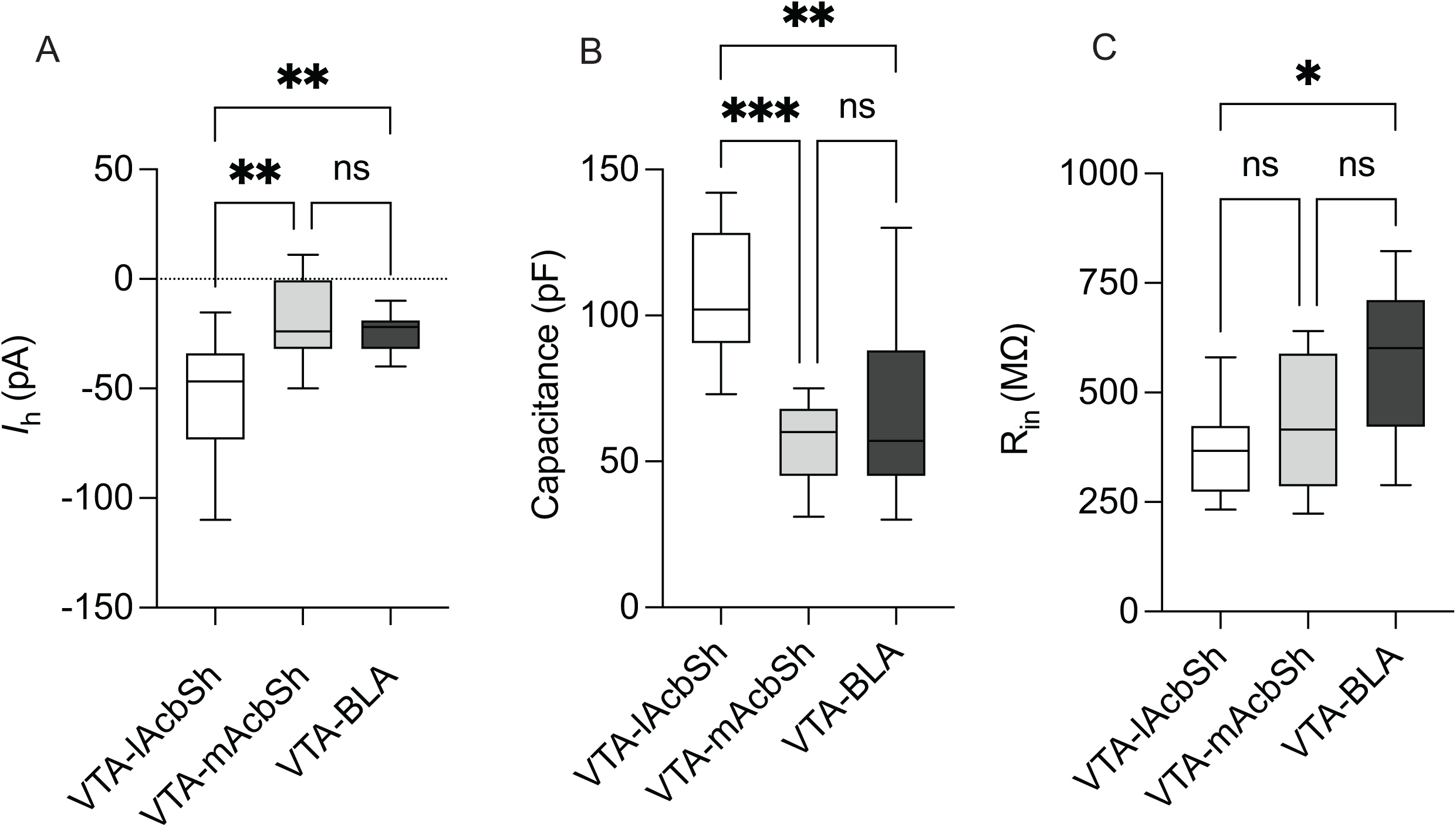
lAcbSh-, mAcbSh-, or BLA-projecting VTA ^DA^ neurons have different intrinsic electrophysiological properties. A) HCN current B) capacitance and C) input resistance of lAcbSh-(open bars), mAcbSh-(shaded bars)- and BLA-(filled bars) projecting VTA^DA^ neurons recorded from ChR2 orexin^cre^ mice.

